# Associations of weather fronts with pig reproduction and mortality

**DOI:** 10.1101/2021.02.01.428991

**Authors:** Tamás Reibling, Linnea Hagstrand, Ákos Maróti-Agóts, Zoltán Barcza, Norbert Solymosi

**Affiliations:** Dunahyb Ltd, Szekszárd, 7100, Hungary; SELAS du Mont-bosc clinique vétérinaire, Bosc-le-hard, 76850, France; Animal Breeding, Nutrition and Laboratory Animal Science Department, University of Veterinary Medicine Budapest, Budapest, 1078, Hungary; Department of Meteorology, Eötvös Loránd University, Budapest, 1117, Hungary; Centre for Bioinformatics, University of Veterinary Medicine Budapest, Budapest, 1078, Hungary

## Abstract

Farmers and practising veterinarians have long suspected the impact of weather fronts on production and animal health. A common impression is that sows will farrow earlier in connection with a cold front. There might be a correlation between daily mortality and the occurrence of a strong atmospheric front. Population-based quantitative studies on weather fronts’ effects on animal health and production are very sparse in the scientific literature. In this study, the associations between the weather fronts and daily farrowing incidence, the pregnancy length and the daily death incidence were analysed. The results show that cold front increased the odds of more than daily six farrowings on the day of the front (with at least 3°C cooling OR: 4.79, 95%CI: 1.08-21.21, p=0.039). On the day of the front, with at least 3°C temperature change both the cold and the warm front increased the odds of the farrowing on the day ≥ 118th day of the gestation (OR: 3.10, 95%CI: 1.04-9.30, p=0.43 and OR: 4.39, 95%CI: 1.73-11.15, p=0.002, respectively). On the day after the day of front, the odds of farrowing on the ≤ 113th day of gestation are increased, if the temperature decrease was at least 2°C the OR: 2.30 (95%CI: 1.04-5.06, p=0.039). On the day after the warm front with at least 1°C temperature increase the odds of more than daily three deaths is increased (OR: 5.44, 95%CI: 1.23-24.05, p=0.025).

## Introduction

In the production animal sector farmers, veterinarians claim weather fronts’ influence on animal health, reproduction and productivity. From the subjective experiences, the practitioners may conclude contrary interpretations of weather front effects on animal health. Population-based quantitative studies on weather fronts’ effects on biological alterations in living beings are very sparse in the scientific literature. The population-based studies may clarify the empirical contradictions. On the other hand, these studies may conclude to contradictory results as well^1^,^2^. Nevertheless, they are more reliable comparing to the subjective impressions. Although there are some papers in the scientific literature publishing quantitative case studies of weather front effects on biological processes^3^,^4^. However, this approach cannot handle the biological variability that should be fundamental to a requirement of the results to be extended to a wider population.

Quantitative studies of the weather front effects are complicated by the debate on weather front definition^5^. The applied approach significantly influences the front identification results, either synoptic (subjective) or automated (objective) methods. As the synoptic approaches have side dependent moments for reproducible front based analyses the usage of automated, objective front identification methods, algorithms may be more justifiable. Besides other methods,^6^ it seems to be recommendable to use of the thermal front parameter (TFP) for automated front identification^7^.

The first author’s experience initiated our study. He is a practising veterinarian, and according to his observations, the cold fronts increase both the daily number of farrowings and pig deaths. The work aimed to analyse quantitatively the associations between fronts identified objectively and some important reproduction, health-related parameters of pig keeping. The parameters used were the daily number of farrowings, pig death and the length of gestation. As we have no found any published quantitative, population-based scientific work on this topic, we think our results may provide an initial point of description of these associations. We hope that if the associations could be identified, further studies would start finding ways to decrease the damage caused by weather fronts on animal production.

## Materials and methods

### Farm data

The pig data was gathered from a farrow to finish farm, in the middle of Hungary (E: 18°47’49.60”, N: 46°29’30.08”).

The farm’s reproduction data has been recorded into a relational database among other production data since the start-up of the farm. All production and reproduction data is joined in the database to sows. In the study, the following reproduction data of sows were used: date of successful insemination and farrowing. Since in the farrowing stall a worker is present continuously, it is possible to record the exact date of the farrowings. The farrowing recorded to a day if it happened between 0 and 24 h on that day. Only those farrowing records were used where the gestation length was between days 112 and 119. This interval was defined as the central 95 percentiles (between 2.5% and 97.5%) of the raw gestation length data. The cleaned data set contains 15,379 farrowing records with the gestation length distribution shown on Fig 1. The farrowing is not synchronised on the farm, and this is important because the synchronisation alternates the timing of farrowing induction fundamentally.

**Figure 1.**
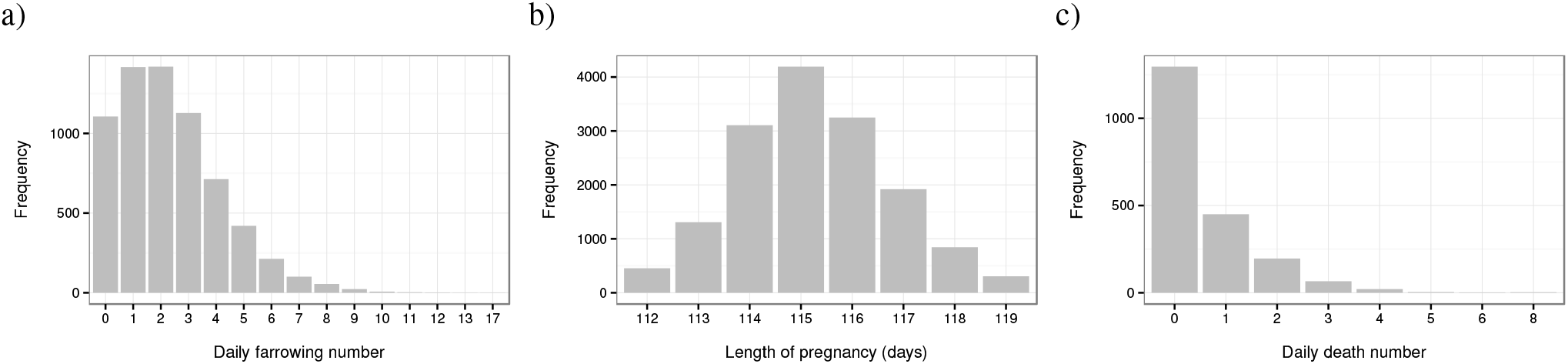
The distribution of daily farrowing numbers (a), gestation length (b) and the daily death numbers (c) in the dataset analyzed

The death data was recorded as a daily number. The numbers were recorded in the following groups; sows, piglets and fattening pigs. Two different methods were used since all units except the farrowing stall are unmanned after 16.00. The death date was recorded for the farrowing sows as the day the animal died since these stables are attended continuously. The keepers leave after 16.00 in the other units, it was recorded as the number of dead pigs found during the morning rounds and the animals who died before 16.00. Consequently, this number may contain animals who died on the certain day and previous afternoon or evening.

The time frame of the analysed data was from 15/09/1995 to 30/11/2013 and 01/01/2009 to 31/08/2014 of the reproduction and death data.

### Objective front

To identify the weather fronts objectively the thermal front parameter (TFP)^8^,^9^ was used. According to earlier studies’ findings^9^,^10^the wet-bulb potential temperature (*θw*) was used to calculate TFP. For masking the threshold −8 K m^−2^ of the TFP was applied as proposed by Berry^10^.

The meteorology data was obtained from the European Centre for Medium Range Weather Forecasts (ECMWF) ERA-Interim reanalysis^11^. For the presented study, the ERA-Interim default spatial resolution was used (0.75° *×* 0.75° grid). For the front identification, the pressure level of 850 hPa, for the near-surface temperature change calculation the 2m level was retrieved from the ECMWF ERA-Interim dataset. According to the availability of pig farrowing and death data, the meteorological data was downloaded for the time period January 1995 to August 2014 at 6-hour intervals.

The front identification was performed for each grid cell, for each day, for each time (00, 06, 12, 18 UTC). A grid cell had front for a certain day if the algorithm identified front in at least three time points in it^6^. Fronts comprising three neighbouring or more cells were used in the analyses^6^.

One of the study’s goals was to find a practically usable front type (magnitude) definition to differentiate the cold and warm fronts we did not used the method of Hewson^9^. Instead of that as Simmonds^12^ proposed the changes between time points in the 2m temperature were used. We defined the temperature changes between two consecutive days by the difference of the minimum temperature of those two days. If the minimal temperature of the day of front was lower than the previous day, then that front was handled as a cold front. If the minimal temperature of the front day was higher than the previous day, then that front was handled as a warm front. The magnitude of the temperature change between the two consecutive days was also used in the study. Grid cell containing the farm geolocation was associated with the farm. If the farm’s grid cell was a part of a front identified, then the farm, the pigs were considered as they are under the influence of the weather front.

### Statistical analysis

To analyse the associations between the biological parameters and the fronts, or the pure temperature changes, logistic regression was used^13^. In the logit models, a dichotomous variable was transformed from the original variables. By this way, the daily farrowing number was transformed by different thresholds to be binary. These thresholds ranged from 0 to 6. The days with the farrowing number above the certain threshold get value 1, the others get value 0 constructing the logit model’s dependent variable.

The values 113-118 (day) were used as thresholds for the dichotomisation for the gestation length. In those analyses, two directions were used to construct a binary variable. If in one direction parturitions occurred, e.g. on the day 113th of the gestation or earlier, then that record got value 1, all other parturitions got value 0. This type of dichotomisations is indicated in the results by, e.g. ≤ 113. In the other direction parturitions occurring, e.g. on the day 116th of gestation or later got value 1, and the other parturitions got value 0. This type of dichotomisations is indicated in the results by, e.g. ≥ 116.

In the daily death number, the method was the same as in the case of the daily farrowing number, but the thresholds ranged between 0 and 3.

The independent variables were the front with temperature changes or temperature changes without front in the logit models. The latter was used to control the importance of the front phenomenon. In the case of independent variables, there were constructed dichotomous variables as well. For example, front days having at least 1°C temperature decrease got value 1, all other days got value 0. In all analyses, the reference (value 0) was all other days not fulfilling the criteria of value 1.

The association between the dependent and independent variables are expressed with odds ratios (OR). ORs quantify how it increases or decreases the odds of any positive outcome in a group compared to other groups. In the present work, the outcome is the binary variable constructed by the approaches described above. The compared groups are defined by the independent variable of the models. All statistical analysis was performed by using the R-environment^**?**^.

## Results

### Daily farrowing frequency

The distribution of the daily farrowing number is shown in Figure 1/a. The median, the mean and the standard deviation of the daily farrowing number is 2, 2.33 and 1.87.

#### On the day of front

The association analysis results between the daily farrowing number and the presence cold front are shown in Figure 2/a. From the plot, one may conclude that odds of daily farrowing number over 6 is significantly higher on days of cold front comparing to all other days. The greater (or equal) than zero°C temperature decrease on the day of the front is associated with doubled odds of more than 6 daily farrowings (OR: 2.35, 95%CI: 1.08-5.13, p=0.032). The greater (or equal) than 1°C, 2°C, 3°C temperature decrease with front has 2.64, 3.97, 4.79 of OR, respectively. This may indicate that the magnitude of temperature decrease (cooling) had a positive association with the odds of the daily farrowing number above six on the day of the front. When we analysed only the temperature change effects without front using the same farrowing number and temperature thresholds, no significant (p<0.05) association was found.

**Figure 2.**
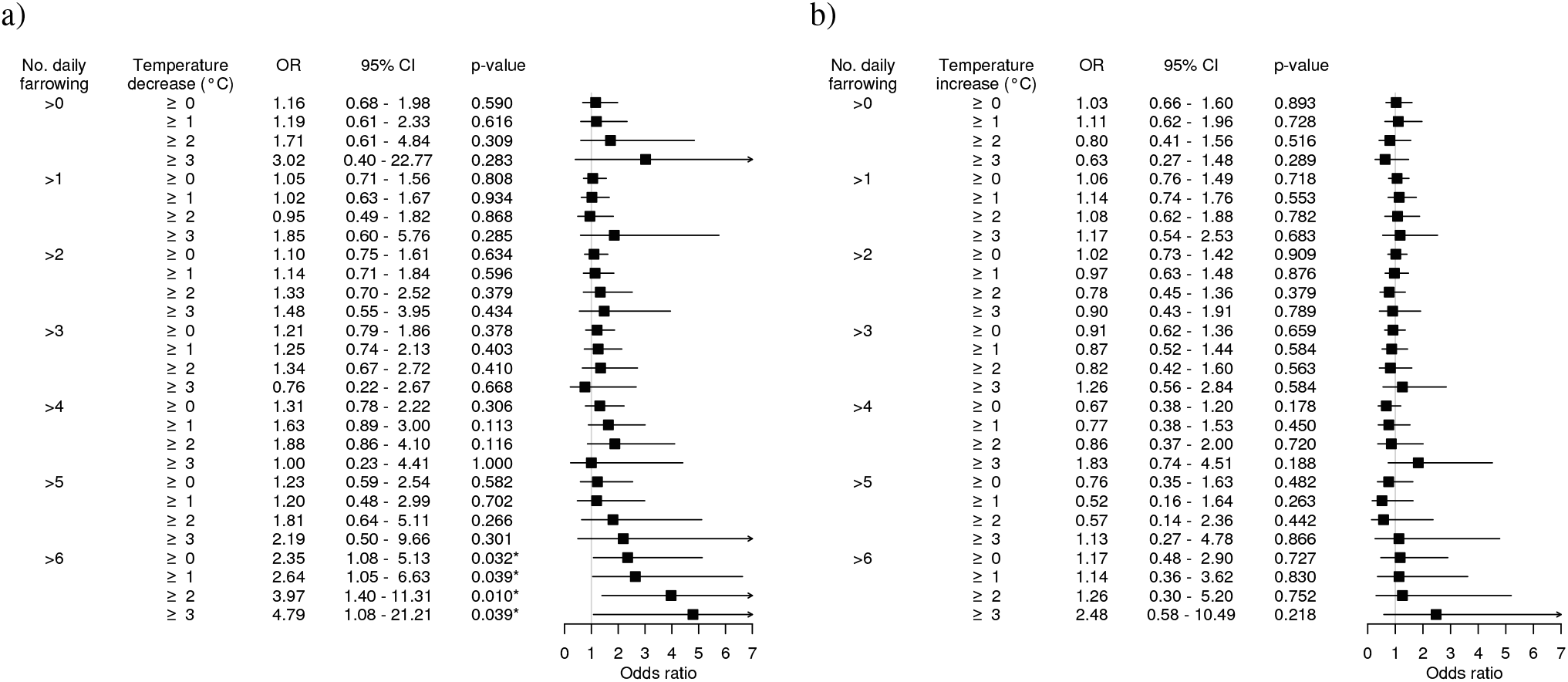
Forest plot (and by side table) presenting the association of cold (a), warm (b) fronts and the daily farrowing numbers. Odds ratios (ORs) express the association between the fronts and the daily farrowing number above a certain threshold (0-6< farrowings/day). The cold and warm property of the front is modelled with the direction and magnitude of temperature between the two consecutive days, based on the 2 meter daily minimum temperatures. On the plot, the black boxes and horizontal lines show the estimates of the ORs and their 95% confidence intervals, respectively. The vertical grey line at value one represents the ineffectiveness. The sign ∗ indicates the significant (p<0.05) association.

Figure 2/b shows the results of the analysis of the link between the daily farrowing frequency and the presence warm fronts. No significant relationship was found. Although the greater (or equal) than 3°C temperature increase shows increased odds of more than six daily farrowings (OR: 2.48, 95%CI: 0.58-10.49, p=0.218), the 95% confidence interval contains the value 1, indicating the result is not significant. As in case of cold front analysing the temperature change affects only (without front information) using the same farrowing number and temperature thresholds there was no significant (p<0.05) association.

#### On day following the day of front

Figure 3/a shows the analysis results between the front and the daily farrowing number on the day following the cold front day. With the same temperature decrease and daily farrowing number thresholds, there was no found significant combination. Analysing the temperature change affects only (without the front information) using the same farrowing number and temperature thresholds there was no significant (p<0.05) association found.

**Figure 3.**
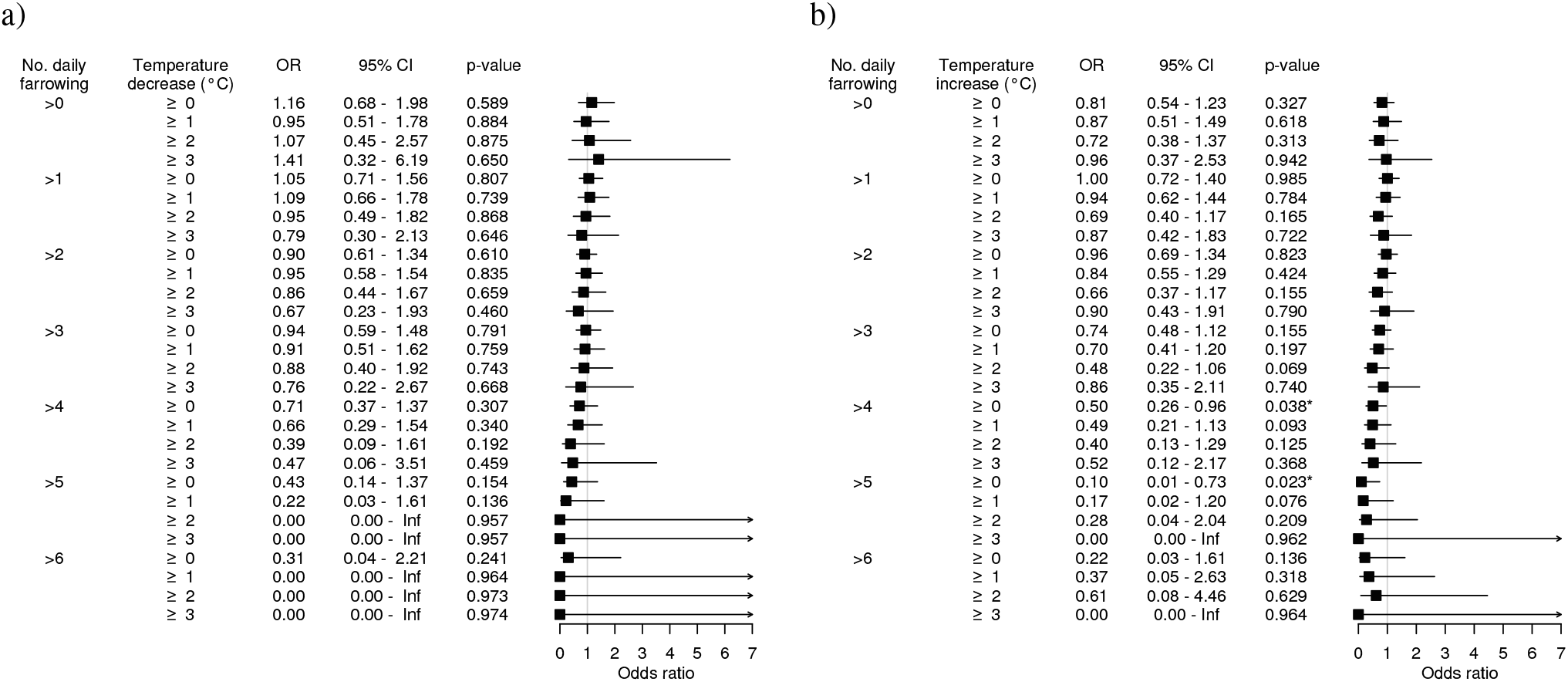
Forest plot (and by side table) of the relationships between cold (a), warm (b) fronts and the daily farrowing number on the day following the day of front.

The statistics describing the association between the front and daily farrowing number on the day following the day of the warm front are shown in Figure 3/b. With most of the temperature increase and daily farrowing number thresholds, there was no found a significant relationship. Two significant threshold combinations show that the warm front may decrease the probability of daily farrowing number above 4 or 5 on the day following the day of the front. The warm front with greater (or equal) than zero °C temperature raising decreased the odds of the daily farrowing number above 4 (OR: 0.50, 95%CI: 0.26-0.96, p=0.038) or 5 (OR: 0.10, 95%CI: 0.01-0.73, p=0.023). Analysing the temperature change affects only (without the front information) using the same farrowing number and temperature thresholds there was no significant (p<0.05) association found.

### Gestation length

The gestation length distribution is shown in Figure 1/b. The median, mean and standard deviation of the gestation length was 115, 115.2, and 1.51 days, respectively.

#### On the day of front

Figure 4/a shows the results of the analysis of the relationship between the presence of cold front and the sow gestation length. According to the figure, it seems the cold front increased the odds of farrowing on later gestation days. On the days of the front when the cooling rate is greater (or equal) than 1°C the odds of farrowing on the day ≥ 117th of gestation is 1.7 times higher than on all other days (OR: 1.74, 95%CI: 1.01-2.98, p=0.045). When on the day of the front the cooling is greater (or equal) than 3°C the odds of farrowing on the days ≥ 118th of gestation is 3.1 times higher than on all other days (OR: 3.10, 95%CI: 1.04-9.30, p=0.043). When the effects of temperature on gestation length was analyzed only (excluding the front information) the temperature decreasing by ≥ 0°C had significant positive relationship with odds of the farrowing on day ≥ 117th of gestation (OR: 0.87, 95%CI: 0.78-0.96, p=0.006). However, the temperature decreasing by ≥ 1°C showed negative relationship with odds of the farrowing on day ≥ 117th or ≥ 118th of gestation (OR: 1.14, 95%CI: 1.01-1.28, p=0.032 or OR: 1.20, 95%CI: 1.04-1.38, p=0.010, respectively).

**Figure 4.**
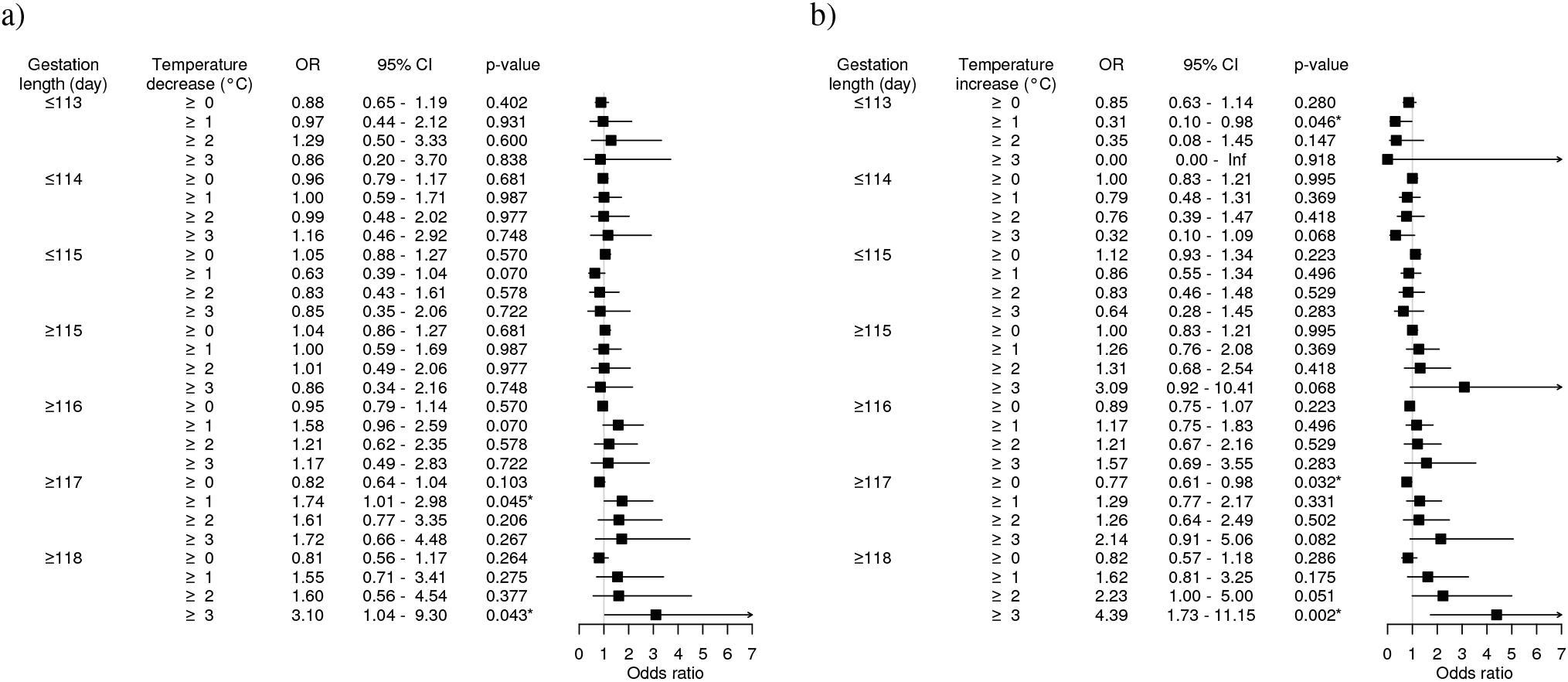
Forest plot (and by side table) of the relationships between cold (a), warm (b) fronts and gestation length.

The analysis results of the relationship between the warm front and the sow gestation length are shown in Figure 4/b. According to the results if on the days of the front the warming is greater (or equal) than 1°C the odds of farrowing on the day ≤ 113th of gestation is 0.3 times lower than on all other days (OR: 0.31, 95%CI: 0.10-0.98, p=0.046). Also, on the days of the front when the warming is greater (or equal) than 0°C the odds of farrowing on the day ≥ 117th of gestation is decreasing, it is 0.8 times lower than on all other days (OR: 0.77, 95%CI: 0.61-0.98, p=0.032). Contrary to these findings on the days of the front when the warming is greater (or equal) than 3°C the odds of farrowing on the day ≥ 118th of gestation is increased, it is 4.4 times higher than on all other days (OR: 4.39, 95%CI: 1.73-11.15, p=0.002). When the temperature effects on gestation length was analyzed without the front information on the days with warming by ≥ 0°C significant changes were found on the odds of the farrowing on day ≤ 115th, ≥ 116th and ≥ 117th of gestation comparing to all other days (OR: 1.11, 95%CI: 1.02-1.20, p=0.018, OR: 0.90, 95%CI: 0.83-0.98, p=0.018 and OR: 0.88, 95%CI: 0.80-0.97, p=0.012, respectively). On the days with warming by ≥ 1°C there were found significant changes on the odds of the farrowing on day ≤ 114th, ≥ 115th and ≥ 117th of gestation comparing to all other days (OR: 1.14, 95%CI: 1.03-1.26, p=0.014, OR: 0.88, 95%CI: 0.79-0.97, p=0.014 and OR: 1.14, 95%CI: 1.01-1.28, p=0.034).

#### On the day following the day of front

The results of analyses of the relationship between the warm front and gestation length farrowing on the day after the day of the front is shown in Figure 5/a. On the figure, one may notice that the temperature decrease by ≥ 2°C increased the odds of farrowing on the day ≤ 113, when the farrowing day is on the day after the day of the front (OR: 2.30, 95%CI: 1.04-5.06, p=0.039). Analysing association between the cooling (without front information) and gestation length farrowing on the day after the day of temperature decrease, no significant relationship was found.

**Figure 5.**
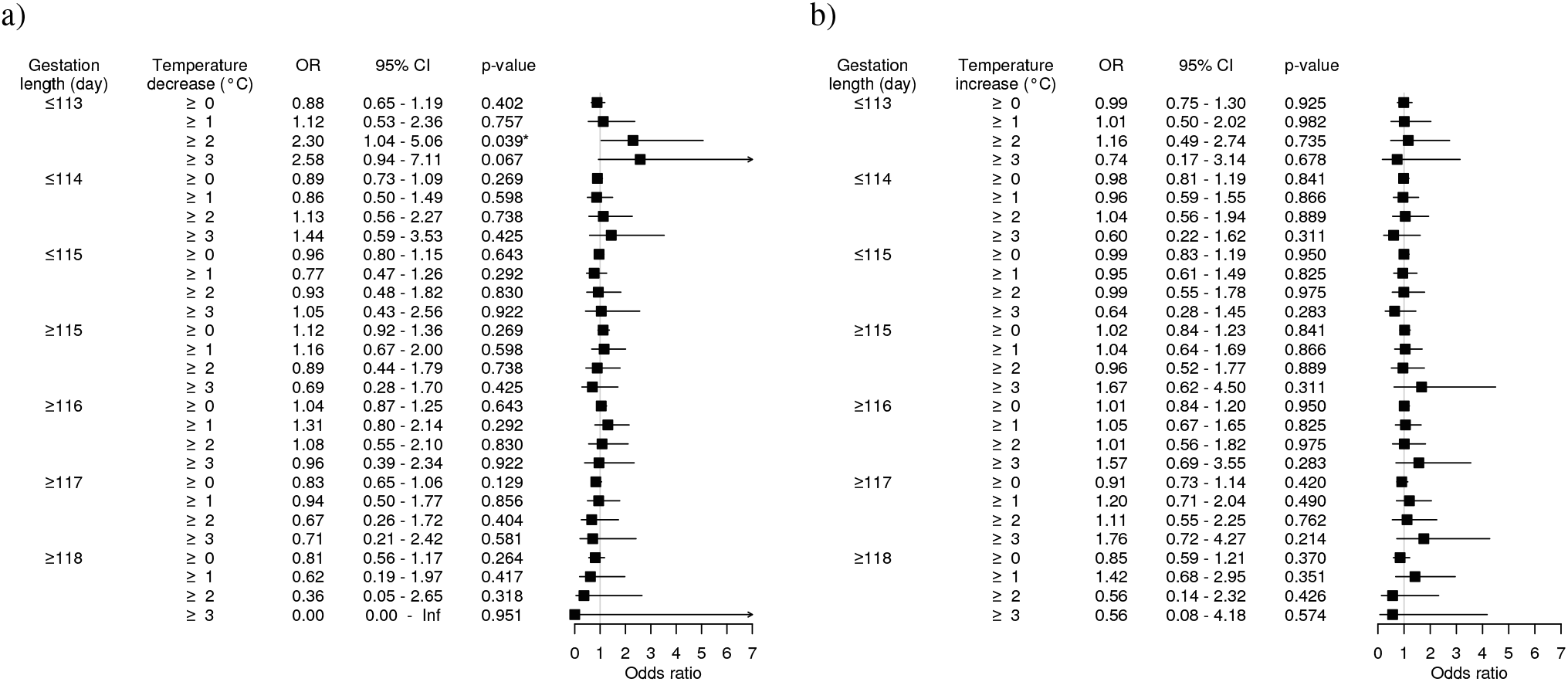
Forest plot (and by side table) of the relationships between cold (A), warm (B) fronts and gestation length farrowing on the day following the day of front.

There was no significant association between the warm front and the gestation length farrowing on the day after the day of the front (Figure 5/b). The temperature change only (excluding the front information) has significant association with the gestation length considering the farrowings on the day after the day of temperature increase. The odds of farrowing on the day ≤ 115th of gestation is greater on the days after the day of ≥ 0°C temperature increase (OR: 1.09, 95%CI: 1.00-1.19, p=0.04). Similarly, the odds of farrowing on the day ≥ 116th of gestation is lower on the days following the day with ≤ 0°C temperature increase (OR: 0.92, 95%CI: 0.84-0.99, p=0.04).

### Death frequency

In the death frequency analyses, we used the daily death numbers of piglets (n=664), fattening pigs (n=425) and sows (n=91) cumulatively (n=1180) and separately as well.

#### On day of front

##### All death

Analysing the death data neither the cold (Fig 6/a) nor the warm (Fig 6/a) fronts had a statistically significant association with the daily death number. Without the front, the temperature decrease had a positive association with the odds of more than 3 daily deaths (OR: 0.44, 95%CI: 0.20-0.96, p=0.039). Conversely, the rising temperature without front effect increased the odds of death. On the days with a temperature increase of at least 1°C the odds of daily death number more than 0 was 1.2 times higher than on other days (OR: 1.24, 95%CI: 1.02-1.49, p=0.029). On the days with temperature increase ≥ 0°C the odds of daily death number more than 3 was 2.3 times higher than on other days (OR: 2.29, 95%CI: 1.04-5.01, p=0.039).

**Figure 6.**
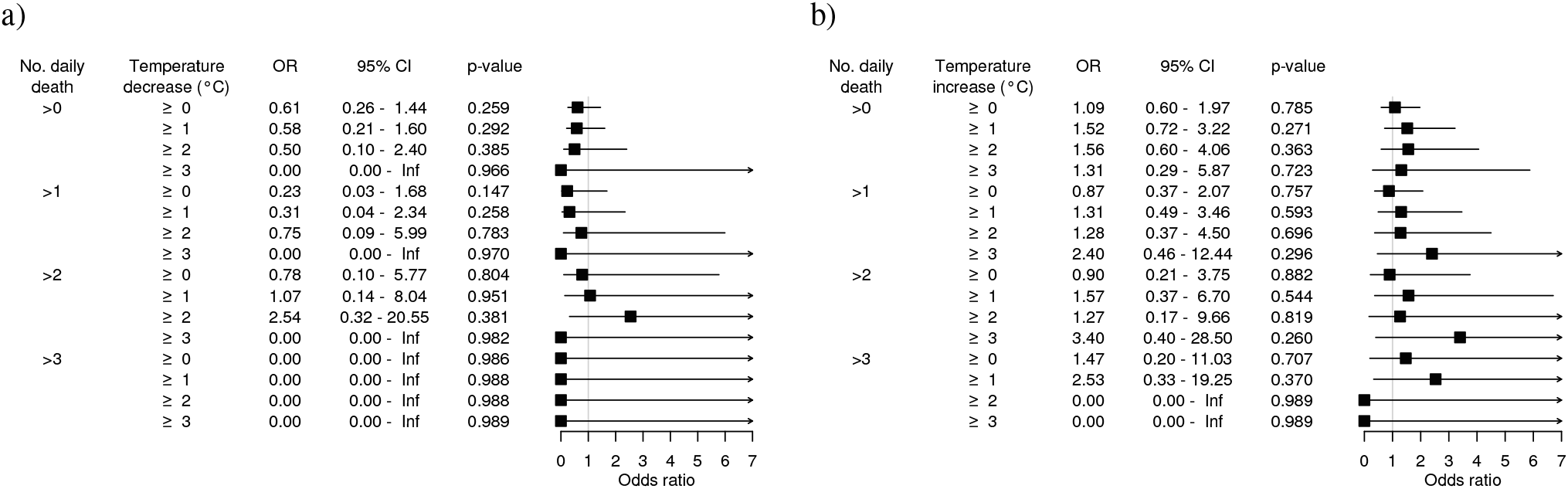
Forest plot (and by side table) of the relationships between cold (A), warm (B) fronts and daily death number on day of front.

##### Piglet death

Neither the cold nor the warm front had a significant association with the daily death frequency. The pure temperature decrease by at least 2°C increased the odds of more than 1 daily death (OR: 1.50, 95%CI: 1.04-2.15, p=0.0291). The temperature increase had no similar effect on the daily death number of the piglets.

##### Fattening pig death

Neither the cold nor the warm front had a significant association with the daily death frequency. Pure temperature decrease had no significant influence on the daily death frequency of fattening pigs. Pure temperature increase by at least 1°C or 2°C increased the odds of more than 0 daily death of fattening pigs (OR: 1.40, 95%CI: 1.11-1.77, p=0.005; OR: 1.34, 95%CI: 1.03-1.75, p=0.032, respectively).

##### Sow death

Neither the cold nor the warm front had a significant association with the daily death frequency. Without a front, neither the decrease nor the temperature increase had a significant association with the daily death number.

#### On the day following the day of front

##### All death

There was no significant relationship found between the cold front and the daily death number on the day after the day of the front (Figure 7/a). No significant relationship was found analysing the temperature only effects on daily death number for the day following the cooling (excluding the front effects).

**Figure 7.**
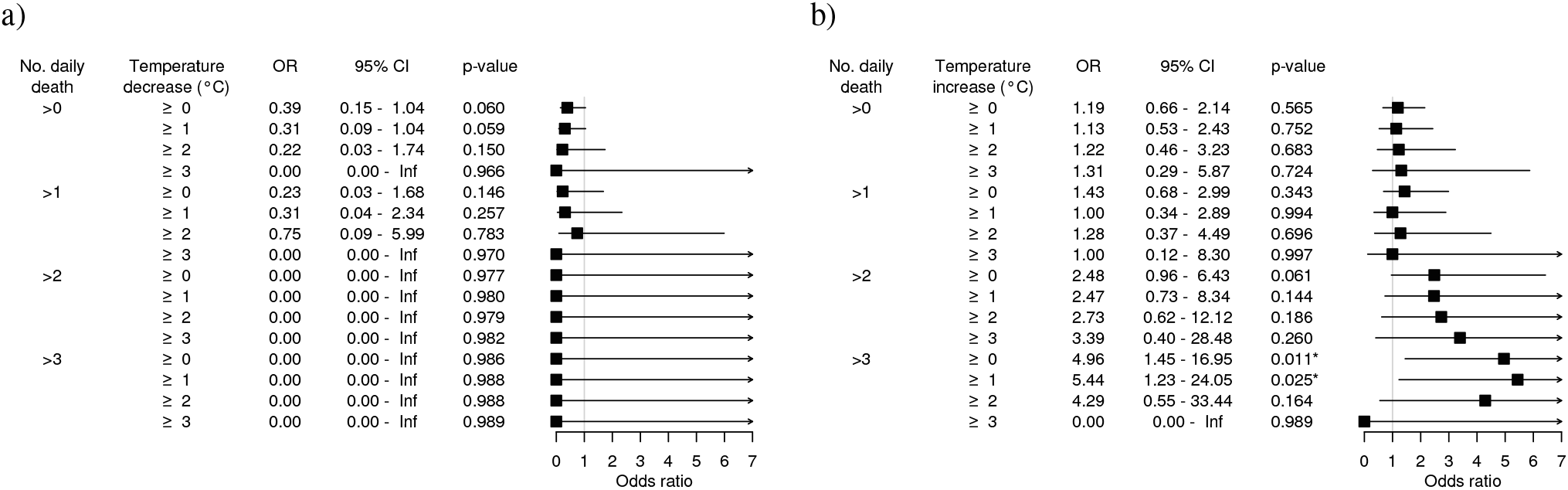
Forest plot (and by side table) of the relationships between cold (A), warm (B) fronts and daily death number on day following the day of front.

The relationships between the warm front and daily death number on the day after the day of the front are shown in Figure 7/b. Warm front with a temperature increase of ≥ 0°C, ≥ 1°C increased the odds of more than 3 daily deaths (OR: 4.96, 95%CI: 1.45-16.95, p=0.011, OR: 5.44, 95%CI: 1.23-24.05, p=0.025). The temperature change only (excluding the front information) had no significant association with the daily death number on the day after the day of temperature increase.

##### Piglet death

Neither the cold nor the warm front had a significant association with the daily death frequency on the day following the day of the front. The results were the same when we modelled the piglet daily death number without the front information, i.e., the temperature changes.

##### Fattening pig death

The cold front had no significant association with the daily death number on the day following the day of the front. On the days following the warm front (with any temperature increase) the odds of daily death number higher than 1 or 2 is increased (OR: 3.71, 95%CI: 1.28-10.75, p=0.016; OR: 11.06, 95%CI: 1.21-100.00, p=0.033, respectively). Without a front, neither the decrease nor the increase of the temperature significantly associated with the daily death frequency on day after day with temperature change.

##### Sow death

Neither the cold nor the warm front had a significant association with the daily death frequency on the day following the day of the front. Without the front, the cooling was associated with decreased odds of death (more than zero daily death). The temperature decrease by at least 0°C, 1°C, 2°C decreased the odds of death on the following day with OR: 0.60 (95%CI: 0.38-0.94, p=0.024), OR: 0.51 (95%CI: 0.30-0.87, p=0.013), OR: 0.48 (95%CI: 0.24-0.93, p=0.029), respectively. Considering the temperature increase without front, if the magnitude of the changes was higher than 0°C, then on the following day there was an increase in the death odds (OR: 1.67, 95%CI: 1.07-2.61, p=0.024).

## Discussion

The cold front significantly increased the odds of more than 6 daily farrowings on the day of the front. This result seems consistent with the practitioners’ impression that the cold front increases the daily farrowing number. This impression may come from the fact that more than daily 6 farrowings can be experienced relatively sparse (Fig 1/a). So if there is a day with such a high number of farrowings it is conspicuous for the farmers, and if they have any information about the presence of weather front, they can easily link the two phenomena.

On the day of the front, the cold front increased the odds of farrowing on the day 117-119th of gestation. On the day of the front, both the cold and warm front increased the farrowing odds on the day 118th, 119th of gestation. However, in the case of the warm front the odds of farrowing on the day ≥ 117 of gestation are decreased. These findings may be interpreted using the following explanation: cold fronts may have a stronger influence on the parturition induction since similar temperature change in the case of cooling increased the odds of farrowing contrary to warming. On the day of the front (if the temperature change is lower) it seems that both the cold and the warm front decreased the odds of the farrowing on the day 112th, 113th of gestation. This association was significant (p=0.046) in the case of warm, but not in the case of the cold front. On the day following the day of the front, the cold front increased the odds of farrowing on the day 112th, 113th of gestation. However, not all thresholds were significant (p<0.05); one may conclude that the magnitude of temperature decrease positively correlates with the effect’s size on the earlier farrowing odds. The links between the weather fronts and the farrowing out of the range 114-116 day gestation length can be interpreted by a supportable argumentation. The sows in the gestation period above the 116th day should be much more prepared for parturition than the sows in the period below the 114th day. This could mean that the farrowing can happen in a shorter time when the sows are in a later period (day 117-119) of gestation. In the case of the sows being in an earlier period (day 112-113), the farrowing may take longer time. Therefore front may cause farrowing on the day of front among the pregnant sows in the later period of gestation. Likewise among the pregnant sows in earlier period front may have protracted effect causing completed farrowing on the day following the day of the front.

On the day following the front, the odds of more than 3 death numbers are increased. The effect size on the odds of death seems correlated with the size of temperature increase positively. Considering the associations of weather fronts and daily death number within the subpopulations, it may be a conclusion that the fattening group dominates this result. For a while, neither among the piglets nor among the sows there is no demonstrable association between the warm front and the daily death number, in the group of fattening pigs the estimates (for daily more than 1 or 2 deaths) are very similar to the estimates for the whole population. Since in the fattening pig population the daily death number is established from death cases of two consecutive days (from 16 o’clock to 16 o’clock) it is not decidable if the front affects on the day of front or the following day. One may speculate that the sudden weather change may affect the animals by perturbating the cardiovascular system control. If this pathogenesis can be accepted, then the death on the day of the front instead of the following day is more plausible. In this case, the result showing the death number increase on the following day of the front might be an artefact due to the cadaver registration protocol.

Some methodological aspects must be detailed to see the limitations of the results correctly. In our study, as reference days were used all days did not satisfy any specified condition. For example, if the cold front day was defined as the day with TFP based front and at least 1°C temperature decrease, then all other days meant the reference or control days. One may propose that the reference days should be selected as days without front, with any non-zero temperature changes. Others may raise that correct reference should be days with TFP based front and at least 0°C temperature increase. Before we decided on the usage of the presented reference, we performed analyses with different control days. All the results came from those models would exceed the limitations of an article. Since the presented results show relatively good concordance with the practitioners’ experiences, we find that this approach can be substantiated.

Another methodological issue is related to the utilisation of 2m temperature change for the front type detection. Day-to-day changes of minimum temperature at the surface are driven by different (but in many cases co-varying) meteorological processes. Fronts are usually associated with clouds with different coverage, which inevitably modulate daily temperature range and minimum temperature. For example, after the overpass of the front clearing skies might cause the lower minimum temperature in the day of front even in case of warm fronts, if the previous day was overcast. Also, in case of cold fronts, clear skies during the previous day might cause lower minimum temperatures than during the day of the front overpass with, e.g. overcast skies. The presence or lack of frontal precipitation might also modulate minimum temperatures. These uncertainties should be kept in mind during the interpretation of the results. Different criteria might be used for the front type identification, but this is out of the present study’s scope. Another important but not exhaustively studied issue is the optimal spatial resolution of meteorological data for TFP based front identification. In the literature, the spatial resolution choice is not supported by scientific reasoning^6^,7,10, it seems to be rather arbitrary. To have deeper inspection into the details of animal health and weather fronts’ relationships, we continue this work in the direction of studying the reference day and spatial resolution issues. Besides these other animal health phenomena (e.g. abomasal displacement in cows, horse colic) in connection with weather fronts are planned to be studied by the similar way.

## Conclusions

By applying the thermal front parameter approach, it was possible to analyse animal health, reproduction, and weather fronts’ associations retrospectively for a long time period. Some results of the analyses are consistent with the empirical impressions of the practitioners in the swine sector. Some other results are difficult to reconcile with everyday experiences. The concordance of the results with the professional experiences suggests that the proposed analytical approach can be recommended for similar weather front related studies. The discrepancies between the results and everyday experiences highlight that some fine-tuning and clarification of limitations of the approach may be required to be extendible for other applications.

